# Antelope space-use and behavioural responses to varying anthropogenic influences may permit long-term persistence in a highly human-dominated landscape

**DOI:** 10.1101/2021.09.15.459901

**Authors:** Rohit Raj Jha, Kavita Isvaran

## Abstract

The only means of conserving a species or a habitat in a human-dominated landscape is through promoting coexistence while minimizing conflict. To achieve this, it is vital to understand how wildlife are impacted by direct and indirect human activities. Such information is relatively rare from areas with high human densities. To investigate how animals respond to altered ecological conditions in human-dominated landscapes, we focussed on a wild herbivore of conservation concern in Krishnasaar Conservation Area (KrCA) in Nepal. Here, blackbuck *Anticope cervicapra*, a generalist grazer, lives in refugia located in a growing human population. We studied the impacts of humans on habitat-use and behaviour of blackbuck. We laid 250 × 250 m grid cells in the entire KrCA and carried out indirect sign surveys with three replications for habitat-use assessment. We observed herds of blackbuck for 89 hours in different possible habitat types, location and time of the day using scan sampling methods. Our habitat-use survey showed that habitats under intensive human use were hardly used by blackbuck, even when high-quality forage was available. In areas with low levels of human activity, natural risk factors, primarily habitat openness, was the major predictor of habitat-use. Interestingly, livestock presence positively influenced habitat-use by blackbuck. Blackbuck were substantially more vigilant when they were in forest than in grassland, again indicating an influence of risk. Overall, blackbuck appear to be sensitive to risk associated with both natural and anthropogenic factors. Our findings have direct implications for managing human-wildlife interactions in this landscape, specifically regarding strategies for livestock grazing in habitats highly used by blackbuck and concerning predictions of how changing land-use will impact long-term persistence of blackbuck. Our work suggests that wild herbivores may be able to persist in landscapes with high human densities so long as there are refuges where levels of human activity are relatively low.

## Introduction

Anthropogenic influences are rapidly pervading habitats worldwide (Maier et al., 2005; Stewart et al., 2010). In the developing world, a large part of the populations or range of many species, including large and charismatic species, occurs within highly human-dominated landscapes. For example, the blue bull *Boselaphus tragocamelus* (Khanal et al., 2018; Meena et al., 2014), blackbuck *Antilope cervicapra* (Jhala & Isvaran, 2016; Krishna et al., 2016), Indian wolf (Sharma et al., 2019; Singh & Kumara, 2006) and the critically endangered Great Indian Bustard, GIB *Ardeotis nigriceps* (Rahmani, 2006; Dutta et al., 2011) are now primarily found among dense human populations in most of their distribution range. A key step towards devising conservation strategies for such species is understanding how they respond to the modifications to their ecology that is brought about by anthropogenic factors.

The major impact of human presence in a habitat is its modification. Human induced modification of landscapes can have adverse impacts on wildlife in the form of habitat loss, fragmentation of already isolated patches, and a continuous increase of direct and indirect human presence (Maier et al., 2005; Stewart et al., 2010). Animals are thought to respond to these modified conditions in different ways. Several studies report shifts in the behaviour of animals in human-dominated landscapes. A study of African lions (*Panthera leo*) in a habitat which had high seasonal fluctuation in wild prey, a stable cattle population nearby, and a high probability of being killed by humans showed that lions mostly preyed upon wild herbivores when available and switched to areas where there are livestock when wild herbivores were low in number. However, even during periods when wild prey were scarce, lions avoided being close to the cattle post, refrained from moving around during times of the day when human movement was maximum, and moved fast when they were in areas with high human movement (Valeix et al., 2012). Riley et al. (2003) and Tigas et al. (2002) also report similar temporal variation in the behaviour of large predators in human dominated landscapes. These studies show that predators refrain from moving around during daylight hours when human movement is maximum. This behavioural modification reduces the time available to locate and catch prey. These studies argue that in the long run, such behavioural changes may compound the human-induced decrease in prey abundance and result in reduced food intake by predators, which in turn can affect their survival and reproduction (Ditchkoff et al., 2006).

Similarly, herbivores also perceive human presence as risk and modify their behaviour accordingly. For example, moose (*Alces alces*) use of sites for browsing depended both on availability of browse and distance from the road indicating that moose trade-off foraging with maintaining distance from the road. The animals even maintained different distances from roads differing in the intensity of human use (Eldegard et al., 2012). Work has shown that when selecting habitat for foraging, herbivores may make complex decisions depending on the amount and type of forage available, associated predation risk, time minimization and energy maximization, state of intra and inter-specific competition, and the influence of direct and indirect human presence (Bjørneraas et al., 2011; Hanley, 1997; Kittle et al., 2008; Krishna et al., 2016).

Given that human activities are known to alter ecological conditions for animal species, understanding how animals respond to these altered ecological conditions is essential for effective conservation planning. For example; GIB, which is endemic to India faces a high risk of extinction with only around 300 individuals existing in eight sub-populations distributed over six states in India (Rahmani, 2006). To conserve the surviving population of GIB, experts have recently recommended engaging with communities in GIB landscapes and forming community reserves where GIB-friendly development activities and traditional land use practices would dominate (Dutta et al., 2011). However, to identify the activities that are GIB friendly, it is crucial to understand how different activities are perceived by GIB. Similarly, most populations of species like blue bull (Khanal et al., 2018; Meena et al., 2014) in Nepal and India; Indian wolf (Sharma et al., 2019; Singh & Kumara, 2006) in India; blackbuck (Jhala & Isvaran, 2016; Krishna et al., 2016) in India live among dense human populations. Approaches to manage and conserve these populations are likely to differ from those used for animals primarily living inside protected areas. We continue to lack key information on behavioural responses of wild animals in landscapes with high human density; such information is required to design conservation strategies for species in these landscapes. Previous work on behavioural responses to anthropogenic factors have primarily been carried out in areas with comparatively low human density (Haidt et al., 2018; Frey et al., 2020; Mendes et al., 2020).

To address how animals respond to altered ecological conditions in human-dominated landscapes, we focused on the behavioural responses of a habitat generalist, blackbuck *Antilope cervicapra*, sharing a landscape with humans in Nepal. This species and study site are well suited to a study of the impacts on anthropogenic factors on the ecology and behaviour of a relatively large, charismatic species of conservation importance. The study site, Krishnasaar Conservation Area (KrCA), is of conservation importance as it is the only landscape which has a wild population of blackbuck in the country. The study species is of high conservation priority in Nepal as there are only around 250 individuals in the wild in the country. The species is locally threatened and is one of 27 mammal species that is legally protected by the Nepal Government under the National Park and Wildlife Conservation Act, 1973 (KrCA, 2017). KrCA represents a conservation area with high human and livestock densities. Merely around 17 sq. km of this habitat is home to around 9000 people, 3000 livestock and around 250 blackbuck.

Blackbuck are appropriate for a study on wild herbivore responses in human-dominated landscapes. They are generalist grazers found in different types of habitats, from semi-dry grasslands to open forest (Ranjithsinh, 1989; Isvaran, 2005). These are open-habitat, group-living animals. They form unstable social groups which change frequently within a single day (Mungall, 1978). These animals are known to use early detection and flight when faced with an approaching risk or predator (Mungall, 1978; Ranjithsinh, 1989). Although currently in most of their range they occur in multi-use landscapes with high human densities, they appear to be risk averse and tend to avoid high levels of human activity (Krishna et al., 2016). Blackbuck face anthropogenic factors commonly experienced by wild herbivores in areas with high human density in the developing world. First, they share foraging areas with livestock, a feature that is common in human-dominated dry landscapes (KrCA, 2017). The impact of livestock on wild herbivores is still not well resolved. Khanal & Chalise (2011) suggests that sharing common foraging space by livestock and blackbuck is not beneficial for the latter as livestock removes a large quantity of resources that could otherwise be used by blackbuck. Also, sharing a landscape with livestock increases the risk of occurrence and transmission of gastrointestinal parasites. Second, blackbuck are frequently reported to feed on crops (Jhala, 1993; Asif & Modse, 2016; Das et al., 2018). Such crop use is one of the main sources of human-wildlife conflict associated with large wild herbivores (Bhatta, 2008; Meena & Jaipal, 2020). While crop fields provide high-quality forage, they are also associated with risks due to direct and indirect human presence (e.g., related to guarding crops). Therefore, in human dominated landscapes there are several factors associated with risks even in the absence of hunting. However, we still lack a comprehensive understanding of how wild species cope with these risks through their use of different habitats and through their social behaviour.

We explored the impact of natural and anthropogenic factors on habitat-use and behaviour patterns by blackbuck in a human dominated landscape. We first focused on habitat use and asked (1) how do blackbuck vary their use of habitats that differ in the level of human activity and in natural ecological conditions like habitat structure and forage abundance? Based on such information available on blackbuck ecology and behaviour, we hypothesized that: a) In human dominated landscapes, resource along with risk factors will jointly affect habitat-use patterns by blackbuck. We expected both natural (closed versus open habitats) and anthropogenic (distance from edge of protected area, livestock densities) risk factors and natural (forage abundance) and anthropogenic (crop availability) resources to affect blackbuck habitat use. We expected blackbuck to balance resource availability against risk factors when making habitat use decisions. b) Livestock are expected to pose both benefits and costs. That is, livestock may remove coarser plant parts and make more nutritious plant parts available to blackbuck but at high densities, may present strong competition for forage (Georgiadis et al., 1989). Therefore, blackbuck should use sites with intermediate livestock densities. Second, we focussed on the behaviour of blackbuck within these habitats and asked (2) what activities do blackbuck carry out in the different habitat types that they visit, and what are the habitat factors that best explain the behaviour patterns in these habitats? Here, we hypothesized that (c) animals would be more vigilant in habitat patches that are perceived as high risk, specifically habitats which are more closed and with higher human activity. (d) Also, as a group-living species, we also expected that animals in smaller herds would be more alert than those in larger herds.

## Materials and Method

### Study Area

We studied blackbuck in Krishnasaar Conservation Area (KrCA), which is situated in Gulariya municipality of western lowland Terai of Nepal (Fig. 1). It lies between 28°7’ and 28°39’N latitude and 81°3’ and 81°4’E longitude. KrCA, measuring 16.95 sq. km. in area, is divided into two areas that experience different management regimes: the Core Area (CA) of 5.27 sq. km and Community Development Zone (CDZ) of 11.68 sq. km. The CA has around 150 households and about 500 cattles. Similarly, the CDZ has 1669 households with a total population of 8789. The total number of livestock recorded from those households was 2384 (KrCA, 2017).

**Fig. 1:**
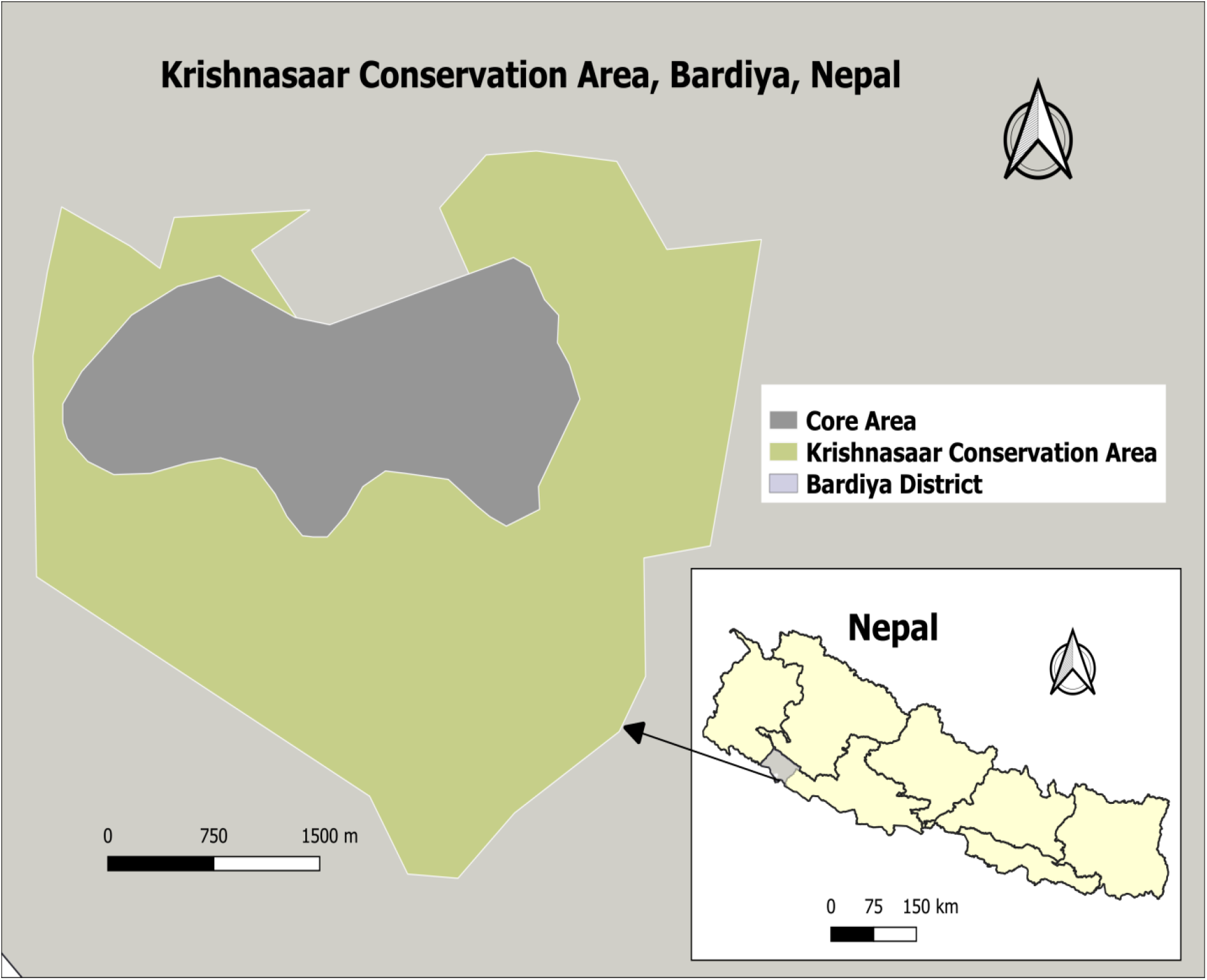
Krishnasaar Conservation Area and its location in Nepal.

### Study species

Blackbuck is a beautiful gazelle-like animal of the family Bovidae in the order Artiodactyla of class Mammalia (Ranjitsinh, 1989), whose natural distribution is limited to the Indian sub-continent (Ranjitsinh, 1989; Khanal & Chalise, 2010). They are medium-sized (30-40 kg), group living and are primarily grazers. The behaviour, nutritional ecology, breeding biology and demography of this species of antelope suggest that they are highly specialized to open, short-grass, semi-arid habitats (Jhala & Isvaran, 2016). Blackbuck continue to persist in such habitats even when there are large human and livestock populations in the area. Thus, this species is suitable for investigating how a species responds to changing ecological conditions in human-dominated landscapes.

### Habitat-use pattern

The study area was divided into 224- 250 m × 250 m grids using QGIS. The sizes of grids were decided based on the daily movement range of blackbuck, which can range from 1.5 km to 5.7 km (Jhala & Isvaran, 2016). This grid size allows us to examine how animals use different habitats available to them. A 20 m buffer was set around the perimeter of each grid and a sampling point was randomly placed within the grid while avoiding the buffer area. This would ensure a distance of at least 40 m between the random points and also ensure independency of sampling points. The main habitat types in these grids were grassland, forest, croplands and builtup areas.

To collect blackbuck sign data as a proxy of habitat use (Krishna et al., 2016), we laid out a strip transect of dimension 20 × 4 m (2 m on either side of the strip) at the chosen sampling point in each of the sub-grids. The strip was divided into five segments of four metres each, and each segment was scored for the presence of indirect signs of blackbuck (pellet groups). Each strip transect was sampled three times over three months (December 2019 to February 2020). This yielded 213 data points from CA and 459 data points from CDZ.

At each strip transect, the four habitat variables, representing potentially important resource and risk factors for blackbuck, were measured; habitat openness (count of number of plants > 1 m in height in a radius of 10 m at the starting point of the strip transect), resource availability (average height and cover percentage of grasses/herbs in a 1 m × 1 m plot measured at 0 m and 20 m points along the length of the strip transect), livestock foraging intensity (number of tracks and dung piles of livestock within the strip transect), and habitat type (grassland, cropland, forest or buildup area).

### Behaviour assessment

To record the behaviour of blackbuck in the different habitat types and zones, we used scan sampling methods for groups (Altmann, 1974). In this method of recording behaviour, a group of animals is selected for observation. The group was defined by including all individuals that are within 50 m of another individual (Clutton-Brock et al., 1982; Isvaran, 2005; Lingle, 2001). We made our observations from raised structures (view-towers) or stood at least 60-80 m distance from the herd. At the start of the observation, we recorded the total number of individuals in the group and their age and sex composition. Age composition was possible only in case of males where the size of the horn and curl in the horn were used to distinguish males as immature male and adult male (Isvaran, 2005). We categorised individuals as fawn (sex not separated), immature male (from horns visible up to three curls in the horn), female (size larger than fawns but no presence of horns), and adult male (more than three curls in the horn). All observations were conducted between 0505 hours and 1815 hours in December, January and February.

An observation session of a group lasted for one hour. Every 10 minutes from the start of the session (zero minute) to the 60^th^ minute, we scanned the group and recorded the behaviour of all individuals at that instant. This gave us seven scans for each hour of observation. These observations were done using a pair of binoculars (Nikon Action EX 8×40 8.2°). During each scan, the activities shown by each individual of different age-classes and sexes were noted. The activities we considered were forage, lie, stand, move, chase, and fight (Meena & Chourasia, 2017) (Supplementary information Appendix 1)). We also noted the type of habitat (grassland, forest, cropland) and the broad zone (CA or CDZ) the group was located in, and weather condition at the time of observation.

### Vigilance behaviour

Vigilance behaviour is a major anti-predator behaviour (Hirschler et al., 2016) that many animals display towards predators and to anthropogenic sources of threat. We examined vigilance behaviour of blackbuck in KrCA to understand whether human related factors influence this behaviour and how blackbuck perceive risk in different habitats (grassland, *Bombax* forest, crop fields, buildup areas) and zones (CA vs CDZ).

We defined a typical vigilance behaviour as when an animal raises its head and scans the surrounding (Beauchamp, 2015). We sampled adult females only to reduce variation in the data that would arise from age and sex related behavioural responses (Isvaran, 2007). To measure vigilance behaviour, we sampled a female from a group continuously for one minute (continuous focal animal sampling, Altmann, 1974). We sampled up to three females from a group. We selected the first female at random, and the subsequent females systematically to ensure that they were at least 20 m away from the first female. If another herd was selected for observation on the same day, it was always located at some distance from the first. During each observation, the number of times the female raised her head up and looked around was noted. The group size, time of the day, habitat type, zone and weather were also recorded for each vigilance observation.

## Analysis

### Habitat-use pattern

For each 20×4 m strip transect, the possible score for habitat use ranged from 0 (no pellet groups of blackbuck) to 5 (pellet groups present in all 5 segments of the transect). Each strip transect was sampled three times over the study period and therefore yielded three scores of habitat use by blackbuck.

#### Resource and risk factors

Each strip transect had two plots where plant height and cover were measured. Plant height and cover thus obtained were multiplied to give an index of resource availability. We averaged these two values to get the average resource available at each strip transect (henceforth, ‘resource abundance’). Livestock indirect signs for each strip transect were summarised as the sum of dung piles and hoof marks of cattle present in the 20×4 m strip. A major risk factor measured for each sampling point (strip transect) was a measure of habitat openness, namely the number of plants over 1 m in height in the 10 m radius plot. Also, the distance of each sample point from the centroid of the core area was calculated. This distance measure was considered as another risk factor because the periphery had settlements, crop fields or forest patches (potential anthropogenic and natural risk factors for blackbuck).

#### Modelling landscape level habitat-use

As blackbuck presence data outside the core area (CA) were scarce (only 2 out of 459 trials), it was not meaningful to examine the predictors of habitat use outside the CA. Therefore, we excluded all the observations outside the core, and used 213 data points obtained from three rounds of sampling inside the CA for modelling habitat use. Since our response variable, the indirect signs score for habitat use, was in the form of count data, we used Generalised Linear Mixed Models (GLMMs) with Poisson error structure. Number of plants over 1 m (index of habitat openness), resource abundance, livestock signs, and distance from centroid of the CA were used as predictor variables. We included two interactions in the model. We modelled the interaction between number of plants over 1 m and resource abundance. Blackbuck are expected to preferentially use more open habitats (i.e., with fewer plants over 1 m). However, in patches with comparatively high resource abundance, blackbuck might respond differently to variation in habitat openness if forage benefits outweigh the costs of risk associated with more closed habitat structure. We also included the interaction between distance from the core centroid and resource abundance, because we expected blackbuck response to the risk factor to depend on resource abundance. The identity of sampling points was treated as a random effect, because sampling points were repeatedly measured. The four predictor variables were checked for multi-collinearity through pair-wise correlations.

All the analyses were run in the software R 4.0.2 (R Core Team, 2013) using the package ‘glmmTMB’ (Brooks et al., 2017). Our statistical inferences were based on the model selection framework using an information theoretic approach (Johnson & Omland, 2004). Based on our hypotheses and knowledge of blackbuck ecology and behaviour, we framed an *a priori* candidate set of 24 models, including both the null and the global models, each representing a different ecological hypothesis (Supplementary information Appendix 2). These models included either single or additive effects of two or more covariates (which included four main effects of the predictor variables and two interaction terms). Models with ΔAIC of <2 were considered to be similarly strongly supported by the data. We used the estimated β-coefficients and their 95% confidence limits to assess the strength of each term in the model. Since no single model appeared to best fit the data, we used multi-model averaging to estimate the parameters using the R package “MuMIn” (Barton K, 2018).

### Behaviour assessment

The data obtained through instantaneous scan sampling were used to calculate the proportion of time animals devote to each activity. Each observation session of a group had seven scans. From these data, the proportion of time spent in a particular activity (e.g., foraging) by an average individual in the group was calculated as the sum of all individuals showing that activity across the 7 scans divided by the sum of all individuals sampled across all seven scans. Thus, for each observation session of a group, the proportion of time spent in foraging, standing, moving, laying, chasing and fighting was calculated. For each session, mean group size across the 7 scans was calculated.

#### Modelling foraging and moving behaviour

We analysed variation among groups in the proportion of time spent foraging, calculated as the total number of records of individuals foraging in a scan sampling session divided by the total number of individuals sampled across all 7 scans in the same session. We used beta regression to fit the model. Beta regression is used when the response variables are probabilities in themselves i.e. the value of the response variable ranges between zero and one (Cribari-Neto F, 2010). Since the response variable had some zeros and ones, violating the assumptions of beta regression, we transformed the proportion data as recommended (Smithson & Verkuilen, 2006). We did not take the approach of adding an offset since some of the values were exactly one.

Similarly, we analysed variation among groups in the proportion of time spent moving, calculated as the total number of records of individuals moving in a scan sampling session divided by the total number of individuals sampled across all 7 scans in the same session. Here too we used beta regressions. The response variable had many zeros and the highest values were well below one. Thus we added an offset to each dependent value to meet the assumption (variable must range from > 0 to < 1) for fitting beta regressions.

The predictor variables for modelling both foraging and moving behaviour were mean group size; habitat type (grassland, *Bombax* forest); location (core and settlement); group type (female only, male only, mixed); weather (no sun, partial sun, sunny); and time of the day (day, evening, morning). The analyses were run using the package betareg (Cribari-Neto F, 2010). We framed an *a priori* candidate set of 26 models for both foraging (Supplementary information Appendix 3) and moving (Supplementary information Appendix 4) behaviour including both the null and the global models, each representing a different ecological hypothesis.

### Vigilance behaviour

To model vigilance behaviour (number of times a female raised her head up in a minute), we used GLMMs with Poisson error distribution. Each female observed constituted a data point.

The predictor variables were group size, habitat type (grassland, *Bombax* forest), location (core, settlement); group type (female only, mixed); and weather (no sun, partial sun, sunny). As multiple females were observed from the same herd, herd identity was treated as a random effect. We framed an *a priori* candidate set of 27 models (Supplementary information Appendix 5) including both the null and the global model.

## Results

### Habitat-use pattern

Out of a total of 672 data points across three months, blackbuck indirect signs were recorded in 99 sampling trials. Of these 97 were from inside the CA and only two from outside. Out of 224 unique sampling points, indirect signs were present in 44 of them. Also, among these 44 sampling points, some were highly populated with signs indicating intensive habitat-use in some of the grids.

Even inside the core area, among the four types of habitats present, no blackbuck indirect sign was obtained from sample points that lay in dense forest. Signs were mostly concentrated in grasslands (67% of 114 trials) and *Bombax* forest (70% of 27 trials), and only one trial (n = 30) in croplands inside the CA showed blackbuck signs (Supplementary information Appendix 6). The habitat use analysis indicated that both risk factors and resources influence blackbuck habitat use (Supplementary information Appendix 7). Blackbuck habitat-use varied with the abundance of plants over 1 m (model averaged weight = 1), resource abundance (model averaged weight = 1) (Fig. 2) and their interaction (model averaged weight = 1). The model averaged coefficients and 95% confidence intervals indicated that when resource abundance was low, habitat use was negatively related to the frequency of tall plants. This relationship weakened as resource abundance increased (Table 1). Habitat-use by cattle also appeared to influence blackbuck habitat-use substantially (model averaged weight = 0.99) (Fig. 3). Blackbuck habitat use was positively related to the use of a habitat by cattle (cattle sign abundance). There was only weak support for the effect of an interaction between distance and resource on blackbuck habitat-use (model averaged weight = 0.52) (Table 1). When resource abundance was low, habitat use was similar at different distances from the centre of the core area. However, in areas with higher levels of resource abundance, habitat use was greatest in the core centre and decreased towards the periphery (Table 1).

**Fig. 2:**
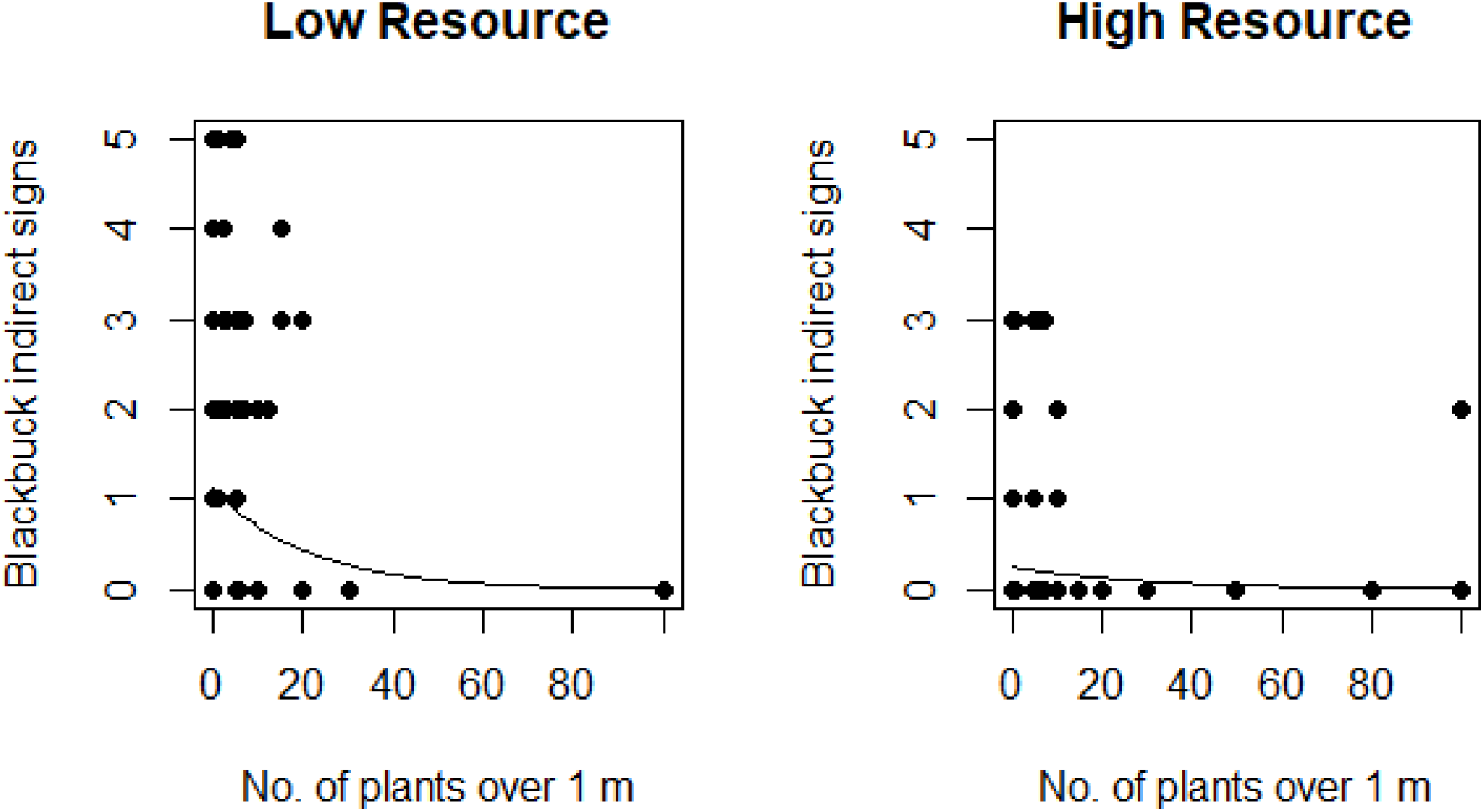
The relationship between blackbuck habitat-use and habitat openness (number of plants over 1 m, a risk factor) at different levels of resource abundance. Resource abundance was modelled as a continuous variable. Here it has been categorised to visualise the statistical interaction between habitat openness and resource abundance. The lines represent model predictions.

**Table 1:** Model averaged β coefficients, 95% confidence limits and weights associated with different predictors in the analysis of blackbuck habitat use (through indirect signs). The model set comprised of 24 models fitted using GLMMs. CL, Confidence Limit; Tall plant abundance, total number of plants over 1 m in height in a plot of 10 m radius laid at the start of the strip transect; Resource, averaged product of plant height and percentage coverage measured in 1 × 1 m plots laid at 1 m and 20 m points on the strip transect; cattle, total number of cattle signs (dung piles and hoof marks) in the strip transect: Distance, linear distance of each sampling point from the centroid of the core. Terms in bold indicate 95% confidence intervals that do not include zero.

|  | **B Estimate** | **95% Lower CL** | **95% Upper CL** | **Weights** |
| --- | --- | --- | --- | --- |
| **Intercept** | **-3.78** | **-5.80** | **-1.77** |  |
| **Tall plant abundance** | **-6.99** | **-10.97** | **-3.02** | **1.00** |
| Resource | 0.005 | -0.71 | 0.71 | 1.00 |
| **Cattle** | **0.22** | **0.09** | **0.35** | **0.99** |
| Distance | -0.02 | -0.41 | 0.37 | 0.78 |
| **Tall plant abundance:Resource** | **2.47** | **1.34** | **3.60** | **1.00** |
| **Distance:Resource** | **-0.48** | **-0.99** | **-0.03** | **0.52** |

CL, Confidence Limit; Tall plant abundance, total number of plants over 1 m in height in a plot of 10 m radius laid at the start of the strip transect; Resource, averaged product of plant height and percentage coverage measured in 1 x 1 m plots laid at 1 m and 20 m points on the strip transect; cattle, total number of cattle signs (dung piles and hoof marks) in the strip transect:

**Fig. 3:**
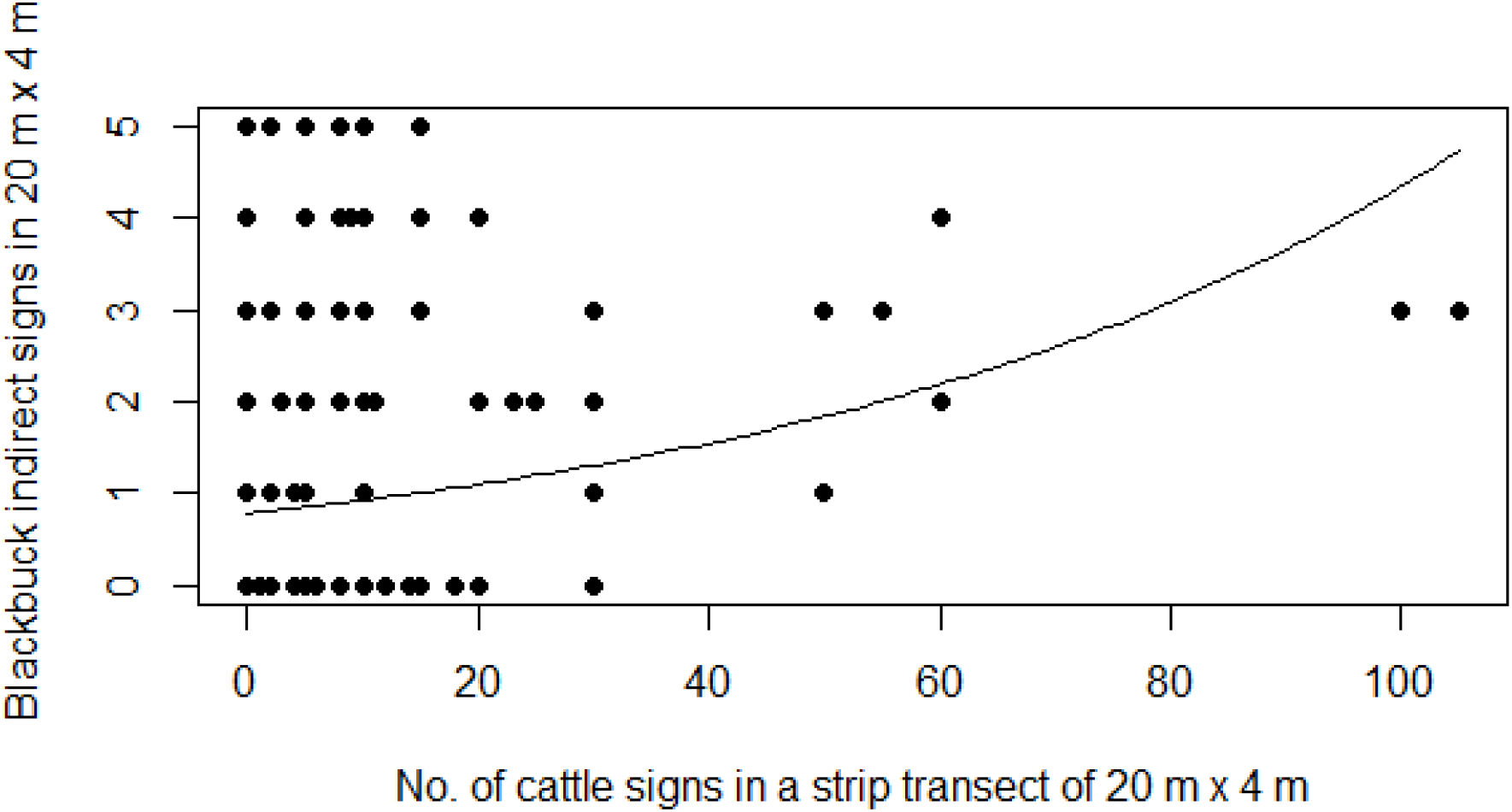
The relationship between blackbuck habitat-use and livestock signs inside the core area. The line represents model predictions.

#### Behaviour Assessment

Although we had planned to observe animals in all possible habitat types, most observations were done in grassland and *Bombax ceiba* forest patches as animals hardly used other habitat patches, at least during the day (also confirmed by the indirect sign data). We scanned animals for 89 hours in total. As each hour (independent observation session) had seven observations, we had altogether scanned 623 groups of animals which comprised 12811 individual instantaneous samples. The mean group size during 89 hours of observation was 20.56 ±1.97 (mean ± SE).

As the youngest animals in the area were already between 8-9 months old (last fawning month was 9 months before the field season started) we could not separate fawn from male and female during observation and a few individuals were unidentified. We did not include unidentified individuals for further analyses. So the analyses included adult males (males with three or more curls in the horn), immature males and females. Animals observed in more than two-thirds of the scans were females.

#### Foraging

Foraging was the most prominent activity observed during scans, forming two-thirds of the total number of scans.

Our model which incorporated only habitat type best predicted foraging behaviour in blackbuck (Supplementary information Appendix 8). Blackbuck foraging behaviour varied most with habitat type (model averaged weight = 0.69). Animals spent a greater proportion of their time foraging in grassland than in forest patch, although this relationship showed some uncertainty (Fig. 4). Other predictors like time of the day (model averaged weight = 0.45), group type (model averaged weight = 0.31), weather (model averaged weight = 0.37), location (model averaged weight = 0.21), and mean group size (model averaged weight = 0.21) did not have much influence on foraging in blackbuck (Supplementary information Appendix 9).

**Fig. 4:**
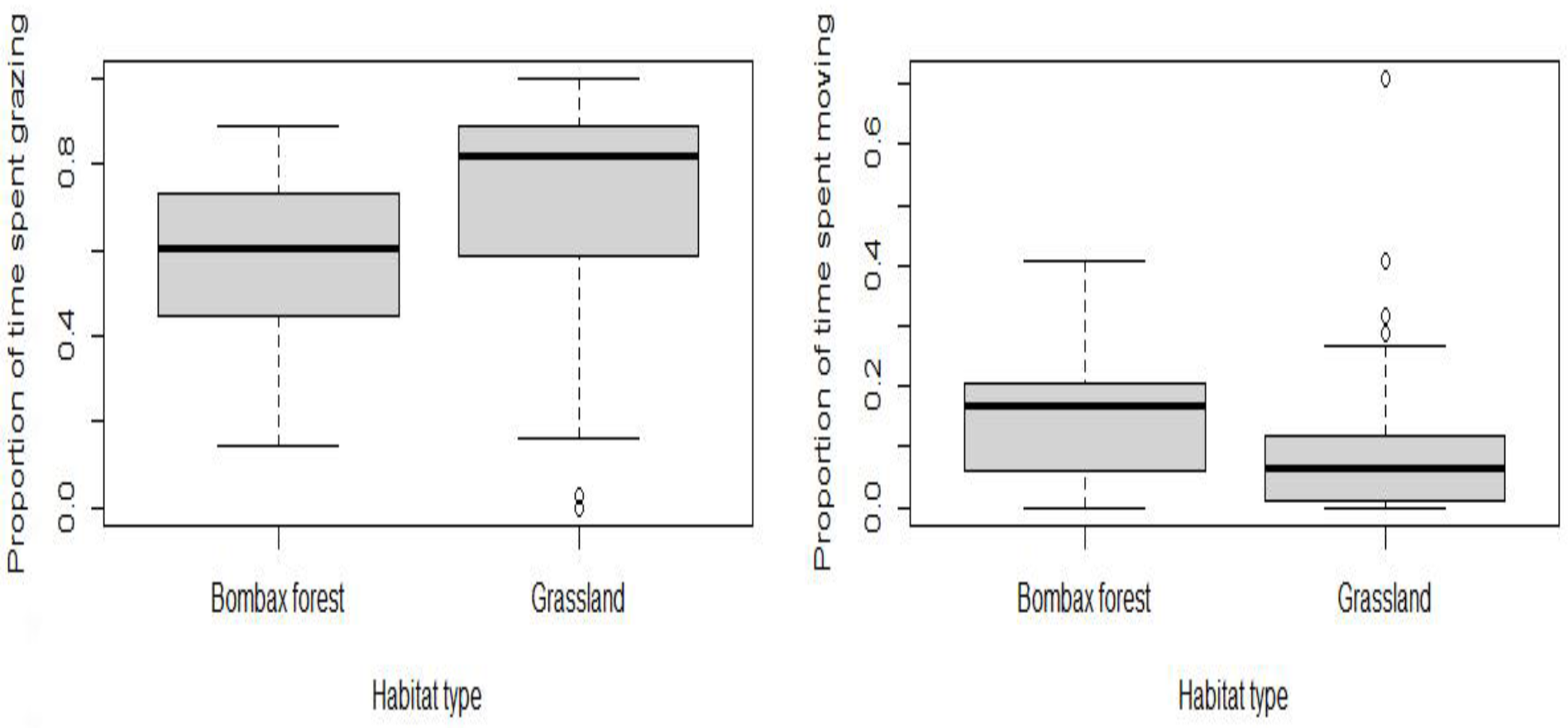
Proportion of time spent foraging (left), moving (right) by blackbuck in two different habitat types. The bold lines represent the median; the box represents inter quartile range and the whiskers represent the data extremes.

#### Moving

Moving was the second most prominent activity observed during scans, after foraging.

Our model which incorporated habitat type, mean group size and time of the day was the best predictor of moving behaviour in blackbuck (Supplementary information Appendix 10). The proportion time spent moving varied most with habitat type (model averaged weight = 0.93). Animals spent less time moving in grassland than in forest patches ( Fig. 4). Other predictors like time of the day (model averaged weight = 0.57), mean group size (model averaged weight = 0.54), location (model averaged weight = 0.19), weather (model averaged weight = 0.05), and group type (model averaged weight = 0.04) did not have much influence on moving behaviour in blackbuck (Supplementary information Annex 11).

### Vigilance behaviour

In total, we observed 186 female blackbuck for one minute each. The mean vigilance frequency was 1.24±0.006 (mean ± SE; range = 0 - 5).

Our model which included group size, habitat type and weather was the best predictor of vigilance by blackbuck (Supplementary information Appendix 12). Vigilance in blackbuck varied the most with habitat type (model averaged weight = 0.93). Females were less vigilant in grasslands than in forest patches. Weather (model averaged weight = 0.84) and group size (model averaged weight = 0.71) were also observed to influence the vigilance behaviour substantially (Fig. 5). Mostly, females were less vigilant on a sunny day than on a completely foggy or partial sunny day whereas vigilance frequency between completely foggy or partial sunny did not differ much. Vigilance decreased as group size increased. There was very little support for the influence of group type (model averaged weight = 0.35) and location (model averaged weight = 0.34) on vigilance behaviour (Supplementary information Appendix 13).

**Fig. 5:**
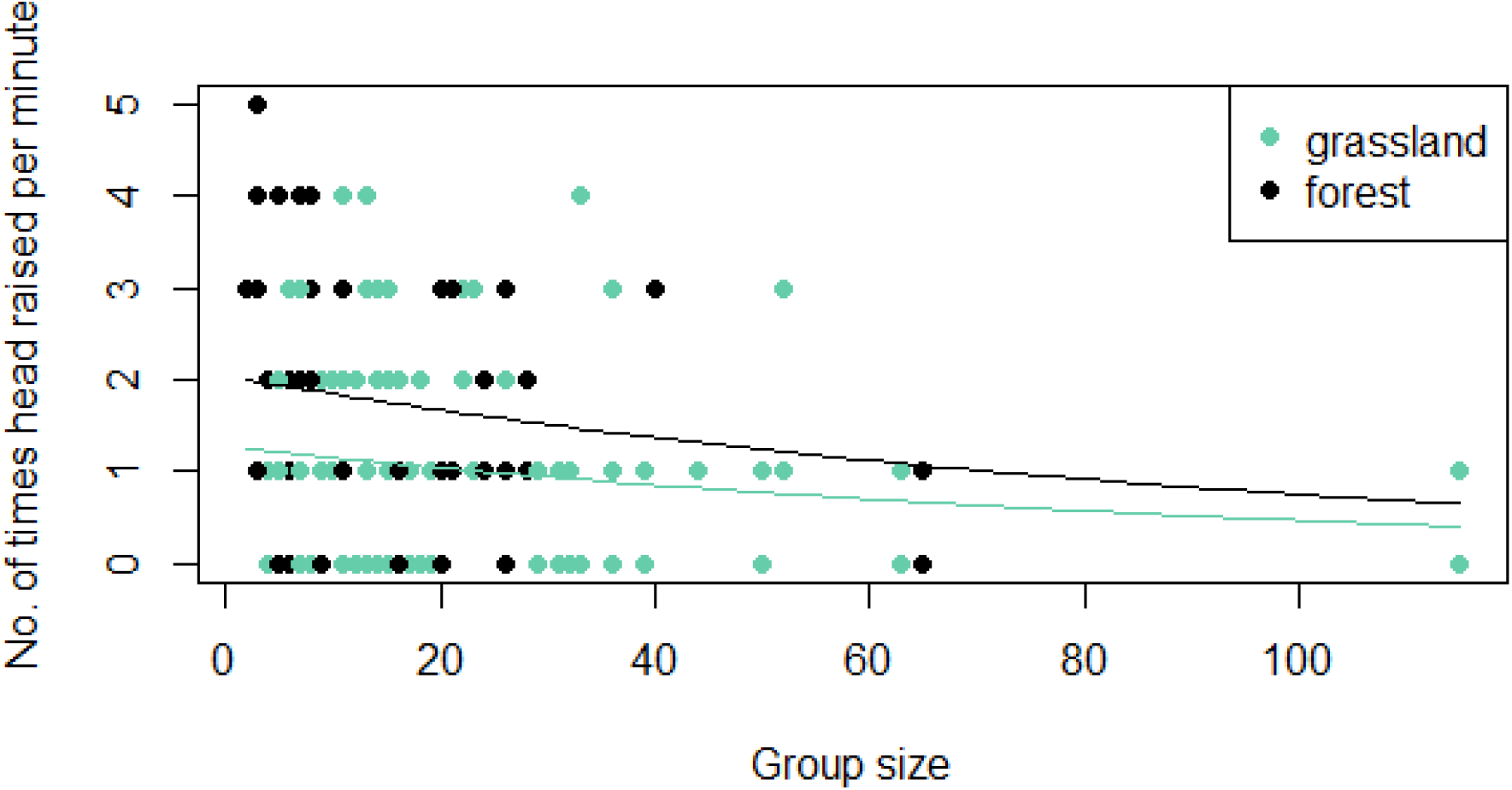
Variation in vigilance frequency in relation to group size and in two different habitat types. Lines are GLMM model predictions using model averaged coefficients.

## Discussion

### Habitat-use pattern

Our findings indicated that habitat use by blackbuck in a human dominated landscape was strongly affected by risk factors, related to both habitat and human activities. First, blackbuck signs in areas outside the core area, in the community development zone (CDZ), were negligible. This indicates a strong negative impact of high levels of human activity on blackbuck habitat use. The CDZ consists of built-up area and crop fields, with no grassland or forest patches. Blackbuck signs were not detected, even in areas of high resource availability within the CDZ, namely crop fields with crops wheat, lentils, and maize. In earlier work based on a social survey conducted at the same site, people have identified these crops as those most used by blackbuck and have reported that these crops were damaged the most by blackbuck (Kuwar, 2015). The lack of blackbuck use of the CDZ suggests that when human activities are high, blackbuck prioritise risk factors over resources when making habitat use decisions. There are several studies that report adverse impacts of human activities on wild ungulate habitat use. A recent study by Costa et al. in 2021shows that both single-season and multi-season farmlands constrict the habitat use of wild ungulates. Similarly, another recent study has found mule deer considerably reducing habitat use in areas with human induced noise (Kleist et al., 2021). These studies, like most previous work, are from landscapes with low densities of humans. Our study contributes to this literature by providing information from a heavily human-dominated landscape.

Interestingly, our study also indicates that blackbuck are able to persist in human dominated landscapes in areas with low levels of anthropogenic factors. Within the core area, blackbuck signs were abundant even though there is some human activity, although at much lower levels when compared with the CDZ. In the CDZ, there were hardly any areas that could not be categorized as either builtup areas or crop fields, whereas such land-uses were limited inside the CA (KrCA, 2017).

Furthermore, within the core area, as we predicted, resource availability and risk factors together best explained habitat-use by blackbuck. Risk associated with habitat, specifically the abundance of tall plants in the area, was consistently negatively correlated with habitat-use by blackbuck. Blackbuck habitat use was the greatest in open habitats without tall plants and decreased sharply as the habitat became more closed. This relationship weakened as resource availability increased. This finding is consistent with studies that have revealed that trade-offs between forage and risk due to predation are fundamental drivers of habitat selection by wild herbivores (Hebblewhite & Merrill, 2009; Krishna et al., 2016). For example, an experimental study in Kruger National Park revealed that large herbivores like African buffalo, wildebeest and giraffe minimized predation risk by avoiding areas with high woody vegetation while foraging (Burkepile et al., 2013).

Blackbuck habitat use did not clearly increase with resource availability. This is likely because blackbuck prefer short grass habitats and are less likely to use tall grass areas. Jhala (1991) found that blackbuck mostly use grassland with short grass with height less than 50 cm and they avoided areas with tall plants. Tall grass can obstruct visibility which is directly associated with predation risk. Furthermore, tall grass is mostly matured with coarse edges and is thus, of comparatively low nutritional quality. Blackbuck selectively feed on more nutritious grass parts and feed less on coarse forage (Jhala, 1997; Jhala, 1991).Krishna et al. (2016) also found that blackbuck prefer relatively short grass areas. This study, which explored blackbuck habitat use in a human-dominated landscape in India, also reported an interaction between resource and risk, as in our study. As another example, in rabbits *Oryctolagus cuniculus*, refuge abundance played a major role in habitat-selection in spring and winter when resources were comparatively abundant, whereas when summer arrived, quality of food available became the sole important factor and the importance of refuge was minimal (Rueda et al., 2008).

Another line of evidence that blackbuck can persist alongside human activity (while levels are low) is provided by our results that blackbuck regularly use areas with livestock presence. This finding has important implications considering the situation where cattle grazing is strictly prohibited in most of the Protected Areas in Nepal and India, the range of blackbuck. This finding is in line with earlier work which has shown that livestock foraging can positively influence wildlife (Schieltz & Rubenstein, 2016). Many recent reviews have also suggested that light to moderate livestock foraging in grasslands is more beneficial in terms of vegetation productivity and quality than a complete lack of foraging (Holechek et al., 2006). This might explain why blackbuck are attracted to those patches of grassland that are used by livestock.

Alternatively, blackbuck might share space with livestock because of resource constraints. In the study area, grazing by livestock along the periphery of the core area, which measures only three sq. km, leaves a very small set of grassland patches towards the centre of the core for blackbuck. Those central grasslands might be insufficient resulting in blackbuck being constrained to forage in areas used by livestock. A previous study has shown that the presence of livestock at fringes of protected areas acts as a buffer between wildlife areas and settlements. However, such a buffer is intact only as long as the resource in wildlife areas is abundant; once the resource depletes, wild animals visit areas used by cattle (Valls-Fox et al., 2018).

Another possible reason for the intensive nature of sharing of foraging space between livestock and blackbuck could be the history of the landscape and the animal itself. One sentence explaining your idea of what might have happened in the study area. A study conducted on translocated wapiti (*Cervus canadensis*) to test the importance of animals being accustomed to an area because of past use has shown that such experiences can impact habitat-use at large and fine scales (Wolf et al., 2009). Further work is needed at our study site to understand the factors influencing the sharing of grazing space between livestock and blackbuck.

### Behavioural variation

Investigating the behaviour of animals in different habitats can provide insights into how animals respond to changing risk and resource factors. We examined behavioural responses likely to be affected by risk. In large herbivores, group living and vigilance are responses to reduce risk from threats such as predation (Isvaran, 2007). Other behaviours, such as foraging and moving, may also provide insights into the costs and benefits associated with different types of habitat.

Vigilance behaviour, a response commonly shown by animals towards threat, as predicted varied with herd size and habitat structure. Animals in larger herds were less vigilant. Such a relationship between vigilance and group size has been shown in many species (Roberts, 1996; Isvaran 2007; Han et al., 2020; Luo et al., 2020). Such a reduction in vigilance with an increase in group size is thought to arise from the benefits of shared vigilance in larger groups. This reduction allows animals to allocate more time to other key activities, such as foraging (Roberts, 1996).

Blackbuck were less vigilant in grassland than in adjoining *Bombax* forest, which had scattered trees with shrubby undergrowth that measured approximately one m in height. Vigilance behaviour is expected to differ with the level of predation risk. As understorey cover obstructs the detection of predators, vigilance rate is expected to increase in such a habitat when compared with open grassland (Ebensperger & Hurtado, 2005). Isvaran (2007) also found that habitat structure played a role in deciding trade-offs between foraging and vigilance. Goitered gazelles were similarly found to change their vigilance rate in response to the level of predation risk (Blank, 2018).

Blackbuck, on average, spent a larger proportion of their time foraging in grasslands than in adjoining *Bombax* forest patches. This difference is likely related to both differences in forage availability and in risk. More undergrowth in forest patches likely reduces grass availability and increases obstruction to vision resulting in increased risk. Like foraging, moving behaviour in blackbuck was also best explained by habitat type. However, in contrast to foraging, blackbuck moved less in grassland and more in *Bombax* forest. This difference might also be explained by the same factors - Foraging and moving behaviour also differed between habitats. obstruction to vision and reduced grass patches due to undergrowth. With the increase in risk and patchiness of grass in *Bombax* forest, blackbuck might move more in this habitat than in grassland.

This study has implications for the long-term persistence of blackbuck in the study site, an area of high conservation priority. First, blackbuck appear to strictly avoid areas largely consisting of agricultural fields and built-up areas and without any grassland or forest patches. At the study site, blackbuck primarily use the core area, which is a relatively small area comprising of grassland and *Bombax* forest within the larger landscape. If direct and indirect human activity increases in the core area too, our study indicates that the long-term persistence of blackbuck in the study area will be negatively affected. Second, KrCA has a history of people grazing their livestock even inside the core area. It continues to be a major demand of people living inside and around the CA even today. A complete ban on livestock grazing is likely to receive opposition from local communities and may thus, pose a challenge to conservation at the study site. As our work shows, livestock foraging does not appear to negatively affect the use of an area by blackbuck. Therefore, management strategies can be explored that permit livestock foraging in selected parts of the core area.

In conclusion, our findings indicate that both ecological and anthropogenic factors influence habitat-use by blackbuck in this human-dominated landscape. Blackbuck appear to be sensitive to risk associated with both natural and anthropogenic factors. Our work suggests that wild herbivores may be able to persist in landscapes with high human densities so long as there are refuges where levels of human activity are relatively low.

## Acknowledgement

We thank the Department of National Park and Wildlife Conservation (DNPWC) of Nepal for research permit (permit dispatch no. 872 dated on 11 November 2019) and National Centre for Biological Sciences (NCBS), Tata Institute for Fundamental Research, Bangalore, India for funding this study. We are grateful to Hariram Yadav for his tireless effort as a field assistant. The corresponding author would like to thank Jayshree Ratnam, Ajith Kumar and Chandni Gurusrikar for administrative support.

## Author Contribution

**Rohit Raj Jha:** Conceptualization (equal); Data curation (lead); Formal analysis (lead); Methodology (equal); Writing-original draft (lead). **Kavita Isvaran:** Conceptualization (equal); Formal analysis (supporting); Methodology (equal); Supervision (lead); Validation (equal); Writing-original draft (supporting).

