## Supplementary information appendix 1 for "Antelope space-use and behavioural responses to varying anthropogenic influences may permit long-term persistence in a highly human-dominated landscape"

### Appendix 1

#### Definitions of terms related to behaviour of blackbuck

| Activities | Definitions |
|---|---|
| Forage | It included both foraging on grass and browse; the individual typically had its head down and was observed cropping grass or browse |
| Move | It referred to the movement of blackbuck which results in change of its location. It included both running and trotting. |
| Stand | It referred to static position of animals when it was on its legs and with its head up. During stand, animals were involved in scanning, socializing or matting activity |
| Lie | Lie refers to position of animals when it is lying on the ground |
| Chase | It referred to condition when an animal or animals followed or went after another animal or animals. All the animals that followed and which were followed were included under chase |
| Fight | It referred to aggressive behaviour showed by animals at each other. All the animals which expressed such behaviour were included under fight. |

#### Appendix 2. Set of candidate models for the analysis of habitat use (indirect sign abundance) by blackbuck

1. Habitat use ~ pover + cattle + resource + distance + pover: resource + distance: resource
2. Habitat use ~ pover
3. Habitat use ~ resource
4. Habitat use ~ pover + resource
5. Habitat use ~ distance
6. Habitat use ~ pover + distance
7. Habitat use ~ resource + distance

8 Habitat use ~ cattle
9 Habitat use ~ cattle + resource
10 Habitat use ~ pover + cattle
11 Habitat use ~ cattle + distance
12 Habitat use ~ pover + cattle + distance
13 Habitat use ~ pover + resource + distance
14 Habitat use ~ pover + cattle + resource
15 Habitat use ~ cattle + resource + distance
16 Habitat use ~ pover + cattle + resource + distance
17 Habitat use ~ pover + resource + pover: resource
18 Habitat use ~ resource + distance + distance: resource
19 Habitat use ~ pover + cattle + resource + pover: resource
20 Habitat use ~ pover + cattle + resource + distance + pover: resource
21 Habitat use ~ pover + resource + distance + distance: resource
22 Habitat use ~ pover + resource + distance + pover: resource
23 Habitat use ~ cattle + resource + distance + distance: resource
24 Habitat use ~ cattle + resource + distance + pover: resource + distance: resource

**Appendix 3. Set of candidate models for the analysis of foraging behaviour by blackbuck**

1 Forage ~ mean group size + habitat type + weather + time + location + group type
2 Forage ~ mean group size
3 Forage ~ habitat type
4 Forage ~ mean group size + habitat type
5 Forage ~ weather
6 Forage ~ mean group size + weather
7 Forage ~ habitat type + weather
8 Forage ~ mean group size + habitat type + weather
9 Forage ~ time
10 Forage ~ mean group size + time
11 Forage ~ habitat type + time
12 Forage ~ weather + time

13 Forage ~ mean group size + weather + time
14 Forage ~ habitat type + weather + time
15 Forage ~ habitat type + group type + time
16 Forage ~ mean group size + habitat type + weather + time
17 Forage ~ mean group size + habitat type + time + location
18 Forage ~ habitat type + weather + time + location
19 Forage ~ mean group size + time + group type
20 Forage ~ habitat type + location
21 Forage ~ weather + location
22 Forage ~ time + location + group type
23 Forage ~ mean group size + time + location
24 Forage ~ mean group size + habitat type + weather + time + location
25 Forage ~ mean group size + habitat type + group type
26 Forage ~ habitat type + time + location + group type

**Appendix 4. Set of candidate models for the analysis of moving behaviour by blackbuck**

1 Move ~ mean group size + habitat type + weather + time + location + group type
2 Move ~ mean group size
3 Move ~ habitat type
4 Move ~ mean group size + habitat type
5 Move ~ weather
6 Move ~ mean group size + weather
7 Move ~ habitat type + weather
8 Move ~ mean group size + habitat type + weather
9 Move ~ time
10 Move ~ mean group size + time
11 Move ~ habitat type + time
12 Move ~ weather + time
13 Move ~ mean group size + weather + time
14 Move ~ mean group size + habitat type + time
15 Move ~ habitat type + group type + time

16 Move ~ mean group size + habitat type + weather + time
17 Move ~ mean group size + habitat type + time + location
18 Move ~ habitat type + weather + time + location
19 Move ~ mean group size + time + group type
20 Move ~ habitat type + location
21 Move ~ weather + location
22 Move ~ time + location + group type
23 Move ~ mean group size + time + location
24 Move ~ mean group size + habitat type + weather + time + location
25 Move ~ mean group size + habitat type + group type
26 Move ~ habitat type + time + location + group type

**Appendix 5. Set of candidate models for the analysis of vigilance behaviour by blackbuck**

1 Headup ~ group size + habitat type + weather + location + group type
2 Headup ~ group size
3 Headup ~ habitat type
4 Headup ~ group size + habitat type
5 Headup ~ weather
6 Headup ~ group size + weather
7 Headup ~ habitat type + weather
8 Headup ~ group size + habitat type + weather
9 Headup ~ location
10 Headup ~ group size + location
11 Headup ~ habitat type + location
12 Headup ~ weather + location
13 Headup ~ group size + habitat type + location
14 Headup ~ group size + weather + location
15 Headup ~ habitat type + weather + location

16 Headup ~ group type

17 Headup ~ group size + group type

18 Headup ~ habitat type + group type

19 Headup ~ weather + group type

20 Headup ~ location + group type

21 Headup ~ group size + location + group type

22 Headup ~ habitat type + location + group type

23 Headup ~ weather + location + group type

24 Headup ~ habitat type + weather + group type

25 Headup ~ habitat type + weather + location + group type

26 Headup ~ group size + habitat type + weather + group type

27 Headup ~ group size + habitat type + weather + location

**Appendix 6: Habitat type-wise distribution of blackbuck indirect signs inside core area**

| S.N. | Habitat type | No. of sample points, summed across three trials | No. of sample points with blackbuck indirect signs | No. of sample points with no blackbuck indirect signs | Percentage of sample points with blackbuck indirect signs |
|---|---|---|---|---|---|
| 1. | Grassland | 114 | 77 | 37 | 67.50 |
| 2. | *Bombax* forest | 27 | 19 | 8 | 70.37 |
| 3. | Dense forest | 42 | 0 | 42 | 0 |
| 4. | Crop fields | 30 | 1 | 29 | 3.33 |
| | **Total** | **213** | **97** | **116** | **45.54** |

**Appendix 7: Top five models (Based on AIC value) that best explain habitat use by blackbuck (See Appendix 2 for details of model parameters)**

| | Models | LogLik | df | AIC | △AIC | Model Weight (w) | Cumulative w |
|---|---|---|---|---|---|---|---|
| 1 | 1 | -257.63 | 8 | 531.98 | 0.00 | 0.65 | 0.65 |
| 2 | 20 | -259.95 | 7 | 534.45 | 2.47 | 0.19 | 0.84 |
| 3 | 19 | -261.20 | 6 | 534.81 | 2.84 | 0.16 | 1.00 |
| 4 | 17 | -266.89 | 5 | 544.08 | 12.10 | 0.00 | 1.00 |
| 5 | 22 | -266.05 | 6 | 544.51 | 12.54 | 0.00 | 1.00 |

**Appendix 8: Top five models (Based on AIC value) that best explain foraging behaviour use by blackbuck (See Appendix 3 for details of model parameters)**

| | Models | LogLik | df | AICc | △AICc | Model Weight (w) | Cumulative w |
|---|---|---|---|---|---|---|---|
| 1 | 3 | 15.01 | 3 | -23.75 | 0.00 | 0.16 | 0.16 |
| 2 | 26 | 21.21 | 9 | -22.14 | 1.60 | 0.07 | 0.23 |
| 3 | 7 | 16.38 | 5 | -22.03 | 1.71 | 0.07 | 0.29 |
| 4 | 15 | 18.65 | 7 | -21.92 | 1.83 | 0.07 | 0.36 |
| 5 | 21 | 16.32 | 5 | -21.91 | 1.84 | 0.07 | 0.43 |

**Appendix 9: Top five models (Based on AIC value) that best explain moving behaviour by blackbuck (See Appendix 4 for details of model parameters)**

| | Models | LogLik | df | AICc | △AICc | Model Weight (w) | Cumulative w |
|---|---|---|---|---|---|---|---|
| 1 | 14 | 106.79 | 6 | -200.55 | 0.00 | 0.24 | 0.26 |
| 2 | 4 | 104.11 | 4 | -199.74 | 0.81 | 0.16 | 0.40 |
| 3 | 3 | 102.97 | 3 | -199.65 | 0.90 | 0.15 | 0.55 |
| 4 | 11 | 105.12 | 5 | -199.51 | 1.03 | 0.14 | 0.69 |
| 5 | 26 | 106.87 | 7 | -198.35 | 2.20 | 0.08 | 0.77 |

**Appendix 10: Top five models (Based on AIC value) that best explain vigilance by blackbuck (See Appendix 5 for details of model parameters)**

| | Models | LogLik | df | AICc | △AICc | Model Weight (w) | Cumulative w |
|---|---|---|---|---|---|---|---|
| 1 | 8 | -254.01 | 6 | 520.50 | 0.00 | 0.21 | 0.21 |
| 2 | 26 | -253.21 | 7 | 521.05 | 0.55 | 0.16 | 0.37 |
| 3 | 27 | -253.53 | 7 | 521.69 | 1.20 | 0.12 | 0.49 |
| 4 | 7 | -255.72 | 5 | 521.77 | 1.27 | 0.11 | 0.60 |
| 5 | 1 | -252.64 | 8 | 522.10 | 1.60 | 0.10 | 0.70 |

**Appendix 11: Model averaged β coefficient, 95% confidence limits and weights associated with predictors of foraging behaviour from a model set comprising 26 models.**

| | **B Estimate** | **95% Lower CL** | **95% Upper CL** | **Weights** |
|---|---|---|---|---|
| Intercept | 0.18 | -0.67 | 1.03 | |
| Mean group size | -0.004 | -0.02 | 0.01 | 0.21 |
| Group type | | | | 0.31 |
| Group type: Male | -0.11 | -0.81 | 0.60 | |
| Group type: Mixed | 0.51 | -0.06 | 1.08 | |
| Time | | | | 0.45 |
| Time: Evening | 0.12 | -0.46 | 0.71 | |
| Time: Morning | -0.44 | -1.2 | 0.32 | |
| Weather | | | | 0.37 |
| Weather: Partial sun | 0.19 | -0.58 | 0.96 | |
| Weather: Sunny | 0.63 | -0.17 | 1.44 | |
| Location | | | | 0.21 |
| Location: Settlement | -0.21 | -0.81 | 0.39 | |

| | | | | |
|---|---|---|---|---|
| Habitat type | | | | 0.69 |
| Habitat type: Grassland | 0.46 | -0.098 | 1.02 | |
| Phi | 2.61 | 1.88 | 3.34 | |

CL, Confidence Limit; Habitat type (Grassland, Bombax forest); Location (Core and Settlement); Group type (Female only, Male only, Mixed); Weather (No sun, Partial sun, Sunny); Time (Day, Evening, Morning). Terms in bold indicate 95% confidence intervals that do not include zero.

**Appendix 12: Model averaged β coefficient, 95% confidence limits and weights associated with predictors of moving behaviour from a model set comprising 26 models.**

| | **B Estimate** | **95% Lower CL** | **95% Upper CL** | **Weights** |
|---|---|---|---|---|
| **Intercept** | **-1.60** | **-2.13** | **-1.07** | |
| Mean group size | 0.0078 | -0.001 | 0.01 | 0.54 |
| Group type | | | | 0.04 |
| Group type: Male | -0.13 | -0.74 | 0.47 | |
| Group type: Mixed | 0.14 | -0.32 | 0.61 | |
| Time | | | | 0.57 |
| Time: Evening | -0.33 | -0.74 | 0.08 | |
| Time: Morning | 0.18 | -0.30 | 0.67 | |
| Weather | | | | 0.05 |
| Weather: Partial sun | -0.05 | -0.69 | 0.59 | |
| Weather: Sunny | -0.001 | -0.67 | 0.66 | |
| Location | | | | 0.19 |
| Location: Settlement | -0.096 | -0.55 | 0.35 | |
| Habitat type | | | | 0.93 |
| **Habitat type: Grassland** | **-0.55** | **-1.03** | **-0.17** | |
| Phi | 10.16 | 7.00 | 13.31 | |

CL, Confidence Limit; Habitat type (Grassland, Bombax forest); Location (Core and Settlement); Group type (Female only, Male only, Mixed); Weather (No sun, Partial sun,

Sunny); Time (Day, Evening, Morning). Terms in bold indicate 95% confidence intervals that do not include zero.

**Appendix 13: Model averaged β coefficient, 95% confidence limits and weights associated with predictors of vigilance behaviour from a model set comprising 27 models.**

| | **B Estimate** | **95% Lower CL** | **95% Upper CL** | **Weights** |
|---|---|---|---|---|
| **Intercept** | **0.71** | **0.09** | **1.34** | |
| **Group size** | **-0.01** | **-0.02** | **0.00** | **0.71** |
| Habitat | | | | 0.93 |
| **Habitat: grassland** | **-0.53** | **-0.91** | **-0.16** | |
| Weather | | | | 0.84 |
| Weather: Partial sun | 0.05 | -0.48 | 0.58 | |
| Weather: Sunny | -0.44 | -0.97 | 0.09 | |
| Group type | | | | 0.35 |
| Group type: Mixed | 0.21 | -0.2 | 0.62 | |
| Location | | | | 0.34 |
| Location: Settlement | 0.16 | -0.2 | 0.52 | |

CL, Confidence Limit; Group size, number of blackbuck in the herd for which female vigilance behaviour was observed; Habitat type (grassland, *Bombax* forest); Location (core and periphery of the core, near settlement); Group type (female only, mixed); Weather (No sun, Partial sun, Sunny). Terms in bold indicate 95% confidence intervals that do not include zero.
